# Exploring TAS2R46 Biomechanics through Molecular Dynamics and Network Analysis

**DOI:** 10.1101/2023.11.21.568072

**Authors:** Marco Cannariato, Riccardo Fanunza, Eric A. Zizzi, Marcello Miceli, Giacomo Di Benedetto, Marco A. Deriu, Lorenzo Pallante

## Abstract

Understanding the intricate interplay between structural features and signal-processing events is crucial for unravelling the mechanisms of biomolecular systems. G protein-coupled receptors (GPCRs), a pervasive protein family in humans, serve a wide spectrum of vital functions. TAS2Rs, a subfamily of GPCRs, play a primary role in recognizing bitter molecules and triggering events leading to the perception of bitterness, a crucial defence mechanism against spoiled or poisonous food. Beyond taste, TAS2Rs function is associated with many diseases as they are expressed in several extra-oral tissues. Since the precise mechanism of TAS2R activation is poorly understood, this work aims to characterize the mechanisms underlying the signal transduction on the recently experimentally solved human TAS2R46 bitter taste receptor using molecular dynamics simulations coupled with network-based analysis. The results show that the allosteric activation of the receptor is associated with more correlated dynamics of the receptor and the formation of an interaction between two helices which mainly convey the signal transferring from the extracellular to the intracellular region. By elucidating the hallmarks of the allosteric network of TAS2R46 under varying conditions (ligand-bound, ligand-free, and transition states), this study has enabled the identification of the unique functional mechanisms of this receptor, thereby establishing a foundation for a more profound characterisation of this intriguing class of receptors.

**Highlights:**

- The dynamics of TAS2R46 in different states were studied through molecular dynamics and network analysis.
- The presence of the bitter agonist increases intra-protein correlations.
- TM3 and TM6 helices mediate the allosteric network in Holo TAS2R46.
- The rotation of Y241^6.48^ residue is pivotal in the allosteric network for the Holo TAS2R46.

## 1 Introduction

The sense of taste, also known as gustation, is crucial for mammals in evaluating the taste and composition of foods (Roper, 2017). Bitter, along with sweet, sour, salty, and umami, is one of the five basic taste modalities and allows to distinguish toxic molecules, providing the last checkpoint before the ingestion of potentially harmful substances. Bitter taste arises from the interaction of organic bitter molecules with type 2 taste receptors (TAS2Rs), which is a subfamily of G protein-coupled receptors (GPCRs) (Chandrashekar et al., 2000). These taste receptors are mainly expressed on the functional gustatory transduction units, i.e. the taste buds of the tongue, which are contained in gustatory papillae. Bitter compounds, binding to TAS2Rs, induce receptor conformational changes and initiate a downstream cascade of events inside the cell typical of GPCR signalling pathways, which ultimately leads to bitter taste perception. It is worth noticing that TAS2Rs are not only located in the taste buds of the tongue, but also several extra-oral tissues express them, such as heart, skeletal and smooth muscle, upper and lower airways, gut, adipose tissue, brain, and immune cells (Behrens and Meyerhof, 2013; Lee et al., 2019). Therefore, their function is not limited to taste evaluation, but it is associated with many diseases, such as asthma or diabetes (Dotson et al., 2008; Liggett, 2014; Pan et al., 2017). Therefore, extra-oral TAS2Rs could also represent a promising target for pharmacological intervention for specific diseases or health conditions. In this scenario, the understanding of the molecular mechanisms driving TAS2R functions is not limited to the taste perception field but can also improve our knowledge of pathologies and relative treatments.

From a structural point of view, TAS2Rs include a short extracellular N-terminus domain, an intracellular C-terminus domain, and seven transmembrane α-helixes (7TMs), which are connected by three extracellular loops (ECLs) and three intracellular loops (ICLs) (Zhang et al., 2017). These receptors present an orthosteric binding pocket which is in the EC part of the 7TMs bundle, involving the extracellular region of TMs II, III, V, VI, VII (Behrens and Meyerhof, 2009; Behrens and Ziegler, 2020; Brockhoff et al., 2010; Pallante et al., 2021). The secondary structure of these receptors is composed mainly of alpha-helix associated with the transmembrane bundle (about 70-75%), whereas there are about 20% of bend, coil, and turn and, in some cases, a minor part of beta-sheets (1-2%), composing the EC and IC domains. The structural similarity between TAS2R receptors and class A GPCRs has in the past led to their classification within the same family. Nevertheless, several recent investigations have pointed out specific characteristics of the bitter receptors, indicating that TAS2Rs can form a distinct family within the GPCRs (Di Pizio et al., 2016; Tokmakova et al., 2023). Indeed, the TM similarity between TAS2Rs and GPCRs is lower than 30% and most of the important conserved motifs of class A GPCRs (DRY motif in TM3, CWxP in TM6, and NPxxY motif in TM7) are missing in TAS2Rs. In TAS2Rs, the DRY motif observed in TM3 of class A GPCRs appears to be substituted by a highly preserved FYxxK motif, while the NPxxY motif in TM7 is replaced by HSxxL (Di Pizio et al., 2016). Moreover, recent literature indicated that TAS2Rs have unique conserved motifs, such as the TM1-2-7 interaction (E^1.42^-R^2.50^-S^7.47^ for TAS2R14), differing from those found in other class A GPCRs, such as the highly conserved N^1.50^-D^2.50^-N^7.49^, with the latter in the NPxxY motif. TM residues are denoted throughout the text using a superscript numbering system based on the Ballesterose-Weinstein (BW) method (Ballesteros and Weinstein, 1995). In this method, the residue corresponding to the most conserved residue in TM X of class A GPCRs is designated as X.50, and subsequent residues are numbered in relation to this position. Moreover, the typical “ionic lock”, which stabilises the inactive state and involves a salt bridge interaction in class A GPCRs, is replaced by a weaker hydrogen bond (HB) between Y^3.50^ and R^6.36^ in TAS2R14. Another important difference is the variation of an important residue related to activation which is W^6.48^ and Y^6.48^ for class A GPCRs and TAS2Rs, respectively (Tokmakova et al., 2023). The comparison between class A GPCRs and TAS2Rs is particularly challenging since no experimental structures of the bitter taste receptors have been determined until recently when cryo-electron microscopy structures of human TAS2R46 in both strychnine-bound and apo states were revealed (Xu et al., 2022). In particular, strychnine is a toxic bitter alkaloid known to be one of the main agonists that activate the TAS2R46-G-protein pathway (Brockhoff et al., 2007).

The understanding of the activation mechanisms of GPCRs is also still unclear. This process is an *allosteric activation* since it transduces various extracellular (EC) stimuli into the intracellular (IC) phenomena: the activation is induced by agonist binding and the subsequent G-protein recruitment. In the last decades, regarding class A GPCRs, a plethora of scientific publications have analysed and discussed GPCR activation mechanisms and allosteric networks (Bhattacharya and Vaidehi, 2014; Bock and Bermudez, 2021; Nivedha et al., 2018; Venkatakrishnan et al., 2016, 2013; Zhou et al., 2019). It is commonly recognised that the outward movement of the transmembrane helix 6 (TM6), which allows binding of the C-terminal part of the G-protein α subunit, is a common feature of GPCR activation triggered by ligand binding in the orthosteric site (Bock and Bermudez, 2021; Venkatakrishnan et al., 2016; Zhou et al., 2019). More in detail, the common activation mechanisms of class A GPCRs from an inactive to an active state involve the elimination of TM3-TM6 contacts, the formation of TM3-TM7 contacts, and the rearrangement of TM5-TM6. Globally, upon binding various agonists, this process triggers the outward movement of the cytoplasmic end of TM6 and the inward movement of TM7 toward TM3 (Nygaard et al., 2009; Rasmussen et al., 2011; Venkatakrishnan et al., 2016; Zhou et al., 2019). TAS2Rs seem to have peculiar features characterising their conformation and allosteric activation, presenting remarkable differences from class A GPCRs. The typical conformational changes related to class A GPCRs were not observed in the recently solved structures, thus leading to the classification of TAS2Rs as class T GPCRs (Xu et al., 2022). Furthermore, the only large conformational change detected was the different localization of the ECL2, which occupied the orthosteric binding pocket in the inactive state. Finally, a rotation of the Y241^6.48^ side chain toward the centre of the 7TMs bundle was observed in the ligand-bound state. Moreover, previous literature suggested that distinctive features of TAS2Rs compared to GPCRs comprise a higher involvement of ECL2 in the binding of agonists and the absence of the ECL2-TM3 disulfide bridge, which is involved in GPCR stabilization. This implies an alternative mechanism for regulating conformational states in TAS2Rs, potentially resulting in a less stabilized inactive state (Di Pizio et al., 2016). Nevertheless, the mechanism underlying the activation of TAS2R receptors remains incompletely elucidated, necessitating further investigation to delineate the distinctive features of these receptors in comparison to class A GPCRs. Moreover, in the absence of molecular structures, it has thus far been infeasible to gain insight into the conformational dynamics of TAS2Rs, thereby hindering an understanding of the principal dynamic molecular mechanisms and structural features underlying receptor function.

In light of the scientific context described above and based on the recent experimental structures of the TAS2R46 receptor (Xu et al., 2022), this study aims to explore the conformational changes and characterise the allosteric networks of this bitter taste receptor induced by the presence or absence of an agonist in the ligand-binding pocket through computational modelling methodologies. Computational techniques are rapidly becoming paramount techniques to highlight the key molecular features linked to the recognition of small molecules by taste receptors, especially for the bitter taste, and identify subsequent molecular events and activation mechanisms (Born et al., 2013; Dagan-Wiener et al., 2019; Di Pizio et al., 2017; Di Pizio and Niv, 2015; DiPizio et al., 2020; Fierro et al., 2022; Levit et al., 2014; Malavolta et al., 2022; Nowak et al., 2018; Pallante et al., 2021). Moreover, the analysis of MD simulations through graph-based approaches is becoming an elective method to study the intra-protein structural communication and crucial residues for protein functions (Cannariato et al., 2023; Fanelli et al., 2016; Melo et al., 2020), as in the case of GPCRs (Bertalan et al., 2020; Bondar, 2022; Siemers et al., 2019). Using MD simulations coupled with network-based techniques, the present study aims to characterize the conformational states of the human TAS2R46 bitter taste receptor and investigate the molecular mechanisms at the basis of the allosteric mechanical communication from the EC to the IC regions. We simulated different receptor conditions, i.e. strychnine-bound, ligand-free, and transition states, to pinpoint major differences in the receptor dynamics and allosteric networks. Consequently, this work, through *in silico* simulations of the dynamic behaviour of the TAS2R46 receptor, offers new insights into the molecular structural mechanisms governing the function of this receptor and establishes the foundation for a comprehensive understanding of the unique functioning features of TAS2Rs.

## 2 Materials and Methods

### 2.1 System setup

The molecular structures of human TAS2R46 were retrieved from the RCSB Protein Data Bank, using the PDB codes 7XP6 and 7XP4 for strychnine-bound and Apo states, respectively (Xu et al., 2022). Only the receptor and the ligand (if present) were preserved from the original PDB files. The missing residues (157-172) of the experimental strychnine-bound TAS2R46 structure were modelled using the corresponding model from the AlphaFold Protein Structure Database (P59540 entry) (Jumper et al., 2021) after root-mean-square fitting on the alpha carbons of the experimental structure. The structure of strychnine was refined using MOE (“Molecular Operating Environment (MOE), 2022.02 Chemical Computing Group ULC, 1010 Sherbooke St. West, Suite #910, Montreal, QC, Canada, H3A 2R7, 2023.,” 2022), predicting the protonation state at neutral pH and salt concentration of 0.15 M. As reported in previous literature (Xu et al., 2022), the tertiary amine of the molecule is protonated and its total charge is +1, as also reported in the DrugBank database for physiological conditions (https://go.drugbank.com/drugs/DB15954).

Therefore, two structures were obtained: (i) the receptor bound to strychnine (Holo) and (ii) in the absence of the ligand (Apo). A third structure, which will be referred to as Trans, was then defined by removing strychnine from the binding pocket of the Holo state.

For each of the three structures, the protein-membrane complex has been built using CHARMM-GUI (Feng et al., 2023) as follows. The homogeneous bilayer membrane was composed of phosphatidylcholine (POPC, 16:0/18:1 acyl chains), as reported in previous literature (Hénin et al., 2006; Zou et al., 2019). The system has been inserted in a rectangular box with dimensions of 8×8×11 nm^3^. This choice allowed us to reach a good compromise between acceptable computational cost and the need to ensure the minimum image convention. Then, the system was solvated by using the TIP3P water model before adding an appropriate number of Na^+^ and Cl^-^ ions to reach a physiological salt concentration of 0.15 M and neutralize the overall system charge. The AMBER19SB force field (Lee et al., 2020; Tian et al., 2020) was used to describe the protein, ions, and water, the Lipid-21 forcefield (Dickson et al., 2022) was used for lipids, and the General Amber Force Field (GAFF2) forcefield (Wang et al., 2004) to obtain the topology for strychnine. System preparation and topology definition were performed directly in CHARMM-GUI as done in previous literature (Born et al., 2013; Pallante et al., 2024, 2023; Sengupta et al., 2018; Zhou et al., 2019; Zou et al., 2019). The protein-membrane complexes in the Holo, Trans, and Apo states are shown in Figure 1.

**Figure 1.**
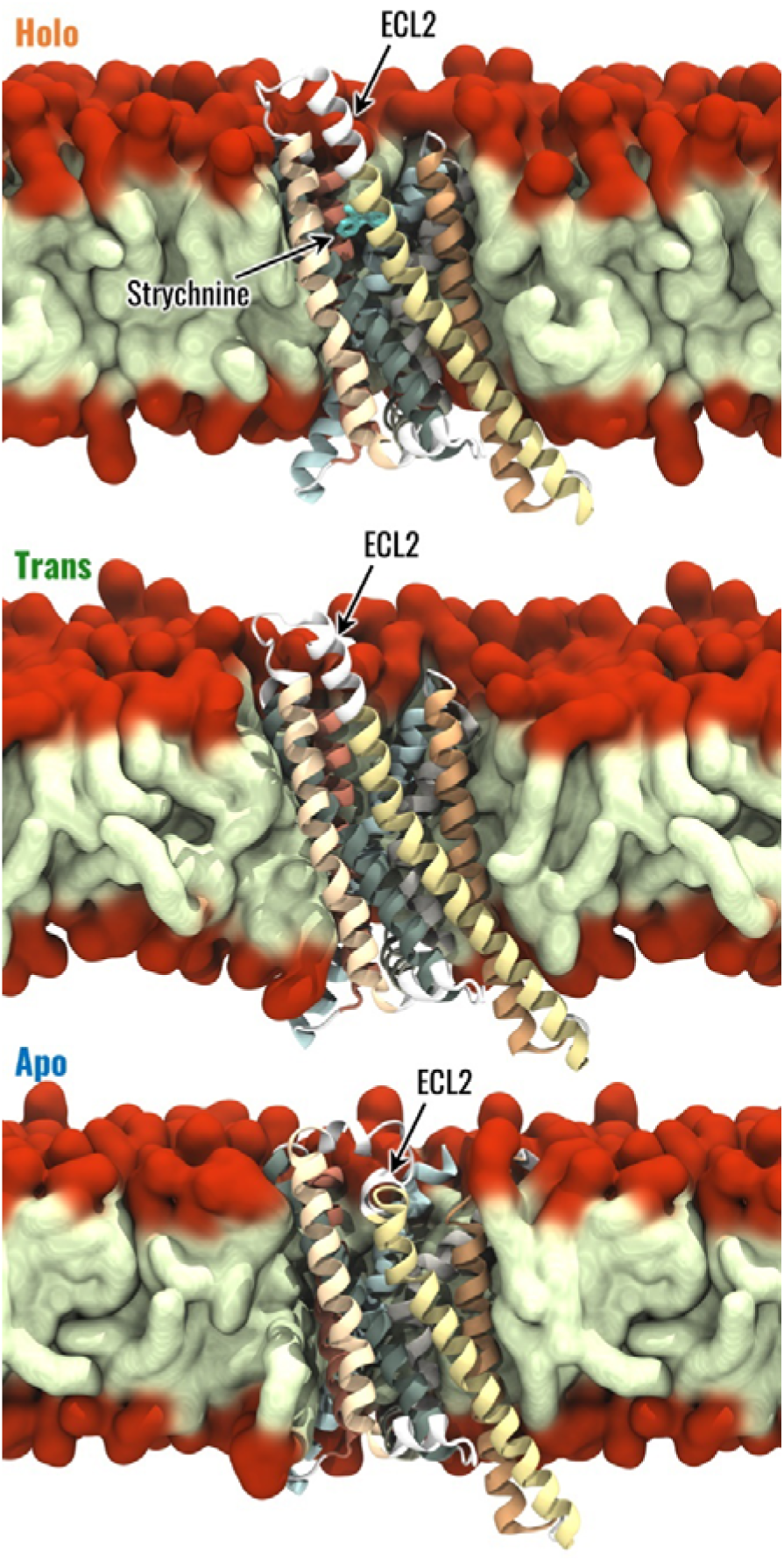
Visual representation of the Holo, Trans and Apo systems. The 7 TMs of TAS2R46 are highlighted with different colours, the hydrophobic tails and hydrophilic head of POPC are coloured in light green and red, respectively.

### 2.2 Molecular Dynamics Simulation

MD simulations were performed using the simulation engine GROMACS 2022 (Bauer et al., 2023) starting from the models described above. For each system, the same simulation protocol was followed as detailed below. First, energy minimization was performed through the steepest descent method for 5000 steps. Then, three replicas were performed following the CHARMM-GUI protocol of equilibration before a production phase of 500 ns long. In detail, six equilibration steps were performed gradually reducing the position restraints on the lipids and protein-heavy atoms (from 1000 to 0 kJmol^-1^nm^-1^ for lipids, from 4000 to 50 kJmol^-1^nm^-1^ for the protein backbone and from 2000 to 0 kJmol^-1^nm^-1^ for protein side-chain heavy atoms). The equilibration phase started with two NVT simulations, performed for 125 ps with a timestep of 1 fs. Then, four NPT equilibration steps were performed, the first one for 125 ps with a timestep of 1 fs, the remaining three for 500 ps with a timestep of 2 fs. The total time of equilibration of each replica was 1.875 ns. The NVT simulations were performed at a reference temperature of T=303,15 K (τ = 1 ps), which is above the phase-transition temperature for POPC, using the Berendsen thermostat (Berendsen et al., 1984), while NPT simulations were carried out at 1.0 bar using the Berendsen barostat with semi-isotropic coupling (τ = 5 ps). Finally, the unrestrained production phase was carried out in the NPT ensemble with a Nose-Hoover thermostat and Parrinello-Rahman barostat for 500 ns. The leapfrog integrator was used, using a time step of 2 fs. The PME algorithm was used for electrostatic interactions with a cut-off of 0.9 nm. A reciprocal grid of 72 x 72 x 96 cells was used with 4th-order B-spline interpolation. A single cut-off of 0.9 nm was used for Van der Waals interactions. LINCS (LINear Constraint Solver) algorithm for h-bonds (Hess et al., 1997) was applied in each simulation step. Three simulation replicas were performed for each investigated system to increase the statistics of the data and ensure the repeatability of the results. Therefore, a total of 4.5 µs of simulation during the unrestrained simulation phase were performed.

### 2.3 Analysis

#### 2.3.1 Conformational analysis

The structural stability throughout the MD simulations for each investigated state (Holo, Trans, Apo) was evaluated by considering the root-mean-squared deviation (RMSD) from the initial configuration of backbone atoms’ positions during the trajectory. To further assess the structural stability of the identified equilibrium, a cluster analysis with the linkage algorithm of the concatenated last 400 ns of each replica was employed, using the RMSD between backbone atoms as metric and 0.15 nm as the cutoff. Moreover, the secondary structure probability for each protein residue was evaluated by considering the concatenated last 400 ns of each replica (Janaszewska et al., 2018). The root-mean-squared fluctuation (RMSF) of alpha carbons was computed in the last 400 ns of each replica, allowing the evaluation of the fluctuations of each residue during the simulation time. Once the structural stability was assessed, the equilibrium trajectories (last 400 ns of simulation) were concatenated obtaining a single 1.2 µs trajectory for each system under investigation. Then, the following analysis was performed considering a sampling time of 50 ps, unless otherwise specified.

The Protein-Ligand Interaction Profiler (PLIP) (Salentin et al., 2015) tool was used to evaluate the specific interactions between strychnine and TAS2R46, underlining the most important residues involved in strychnine binding and the types of interactions established (i.e. hydrogen bonds, hydrophobic interactions, salt bridges, etc.). In detail, the probability of a specific interaction between the receptor and strychnine has been evaluated by considering the interactions on each frame and then averaging the number of occurrences of the interactions on the total number of frames, as done previously (Miceli et al., 2022).

The binding pocket volume was evaluated using the Epock tool (Laurent et al., 2015). The maximum englobing region (MER), i.e., the region of space delimiting the binding pocket, was defined as a sphere of radius 1.3 nm located at the centre of mass of strychnine, considering the starting configuration of Holo replicas. Before the binding pocket volume evaluation, the concatenated trajectories were RMS fitted on the configuration used to define the MER. Moreover, in the analysis, the residues 153 to 176 and, in the Holo system, strychnine were not considered to allow a better comparison between the three systems under study.

Then, the conformation of Y241^6.48^, whose side chain is characterized by a different position in the experimental structures of human TAS2R46, was described by the angle θ, defined as the angle between the centre of mass of Y241 aromatic ring, the alpha carbon of Y241, and the alpha carbon of Y271^7.45^.

The calculation of RMSD, RMSF, and cluster analysis was performed using GROMACS, whereas the secondary structure was evaluated through the STRIDE (Heinig and Frishman, 2004) software package. Conformational analysis was performed through custom-made scripts in Python using the MDAnalysis module. All plots were generated using the Matplotlib (Hunter, 2007) and Seaborn (Waskom, 2021) libraries, whereas the three-dimensional representations of receptor structures were rendered in Visual Molecular Dynamics (VMD) software (Humphrey et al., 1996).

#### 2.3.2 Generalized correlation analysis

A correlation analysis was also employed to identify the regions of the receptor more correlated with each other in the Holo, Trans, and Apo states. In particular, the generalized correlation coefficient (*r_MI_*) was computed as it takes into account linear and non-linear contributions to correlations (Lange and Grubmüller, 2005). This analysis was performed since correlated motions are essential for the biomolecular function of several systems, such as orthosteric and structural signal transduction in GPCRs (Scheer and Cotecchia, 1997).

The generalized correlation coefficient between residues *i* and *j* has been computed as:

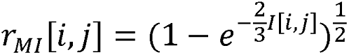

where *I*[*i,j*] is the mutual information between the positions of residues *i* and *j*, where the position of one residue was defined as the position of its alpha carbon. The mutual information has been computed using the density estimator described by Kraskov et al. (Kraskov et al., 2004) with neighbour parameter *k* of 6 as done in previous literature (Lange and Grubmüller, 2005; Melo et al., 2020):

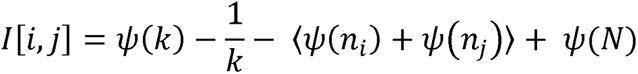

where *N* is the total number of simulation frames, ψ*(x)* is the digamma function, ni is the number of frames in which residue *i* is close to the one in the reference, and ⟨ ⟩ stands for the average by where *N* is the total number of simulation frames, ψ*(x)* is the digamma function, n_i_ is the number of varying the reference frame over the trajectory. For the generalized correlation coefficient calculation, a trajectory sampling time of 1 ns was considered to avoid considering correlated frames in the analysis as done in previous literature (Cannariato et al., 2023; Manrique et al., 2023).

#### 2.3.3 Dynamic Network Analysis

To investigate structural communication within the TAS2R46 receptor, the MD simulations were analyzed through the Dynamical Network Analysis approach, using the dynetan library (Melo et al., 2020). This specific network analysis was selected as it is based on the generalized correlation coefficient, thus considering nonlinear contributions to amino acid dynamical correlations. In detail, in the Dynamical Network Analysis, each protein residue was represented by a node located in its alpha carbon and strychnine was modelled through a single node in the closest atom to its centre of mass. Nodes were linked with edges if the frequency of contacts between the corresponding residues was greater or equal to 0.75, considering two residues to be in contact at a simulation frame if the shortest distance between their heavy atoms is lower than 0.45 nm. The edges of the obtained graph were weighted using the generalized correlation coefficient (Lange and Grubmüller, 2005), computed using a sampling time of 1 ns (Manrique et al., 2023). The network was characterized in terms of betweenness centrality and eigenvector centrality. The betweenness centrality of an edge or node is described as the fraction of the shortest paths in which the considered edge or node is involved and highlights the importance of edges and nodes for the connection of distant parts of the network. In this study, the shortest path between two nodes was defined as the path that maximizes the sum of correlations between the nodes involved in the path. The betweenness centrality is computed as:

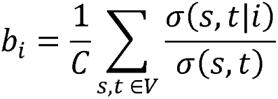

where *σ*(*s, t*) is the number of shortest paths between nodes *s* and *t*(*s, t*|*e*), is the number of such paths passing through node *e*, *V* is the ensemble of graph nodes, and *C* is a normalization factor to allow the comparison of networks with different numbers of nodes. In particular, for a graph of *n* nodes, is equal to

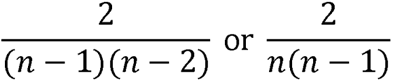

for node and edge betweennesses, respectively.

The eigenvector centrality of a network measures the influence of a node in a graph considering the topology of the graph itself, such that the centrality of a node depends on the centrality of its neighbours. Therefore, if a node has many connections with nodes of small influence, it will also have a small centrality in the network. On the other hand, if a node has few connections with nodes very influential in the network, it might have high centrality because of its indirect influence within the graph. Mathematically, the eigenvector centrality of node *i* is the *i-th* entry of the eigenvector (*x*) of the adjacency matrix weighted on the correlation between the nodes (*A*):

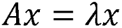

## 3 Results

From the visual inspection of the RMSD plots, it was concluded that the structural stability of the systems on each replica was reached after the first 100 ns (Figure S1). The structural stability was confirmed by the cluster analysis of the concatenated trajectories (last 400 ns of each replica), which showed only one cluster per system. Moreover, the stability of the receptor structure was analyzed in terms of secondary structure probability. This analysis highlighted that the TMs maintained their structure across the simulations without any remarkable alteration of their secondary structure regardless of the receptor’s configuration state (Figure S2). Once the structural stability was confirmed, the Holo, Trans, and Apo systems were characterized as follows. First, we analyzed the difference between the systems in terms of the conformation of the receptor. Subsequently, we examined the dynamic correlations within the structure of the receptor to emphasize distinctions and similarities in the dynamic properties contingent on the TAS2R46 state. Finally, we considered how the difference in conformation and correlation result in different patterns of structural communication inside the receptor through the dynamic network analysis.

### 3.1 Conformational features of TAS2R46 activation

This paragraph describes the conformational analysis conducted to identify remarkable differences among the three analyzed states - Holo, Trans, and Apo. The objective of these analyses is to identify specific structural characteristics associated with the different states of TAS2R46.

The Holo, Trans, and Apo states were initially characterized in terms of RMSF, which was computed to evaluate the flexibility of the different receptor regions. As expected, the most flexible regions of TAS2R46 were the unstructured ones and some differences have been observed between the three receptor states for ICL3 and ECL3 (Figure S3). The Trans state is characterized by higher fluctuations in the ECL3, but the flexibility of the ICL3 in this state is the same as the Holo state. Interestingly, the ICL3 region is more stable and less flexible in the Apo state compared to the other two. On the other side, the ECL2 displays similar fluctuations in the three systems, although it is characterized by a different conformation in the Apo state compared to the Holo and Trans states (Figure 1). Finally, the interaction between strychnine and the receptor was analyzed in terms of type and stability using PLIP. This analysis revealed three main interactions with a probability greater than 0.5, namely two hydrophobic interactions with residues Y85^3.29^ and W88^3.32^ and a salt bridge interaction with E265^7.39^ (Figure S4). This confirms that the main interactions detected in the experimental structure are conserved and involve the TM3 and TM7, highlighting the stability of the ligand inside the receptor’s binding pocket during the simulations.

Since a rotation of the Y241^6.48^ side chain from pointing outward the 7TMs bundle to pointing into the core of the receptor has been previously reported for TAS2R46 (Xu et al., 2022), the conformation of the Y241^6.48^ side chain has been defined in terms of the angle θ, shown in Figure 2A, defined as the angle between the centre of mass of Y241^6.48^ aromatic ring, the alpha carbon of Y241^6.48^, and the alpha carbon of Y271^7.45^. According to its definition, higher values of θ can be related to conformations in which Y241^6.48^ points towards the centre of the receptor. The results show that the conformations of Y241^6.48^ during the MD simulations remained in line with the experimental observations, with higher θ values for the Holo state. Interestingly, the removal of Strychine seemed to affect Y241^6.48^ conformation, whose distribution of θ in the Trans state was located between the ones of the Holo and Apo states. However, the different conformations of Y241^6.48^ are not related to outward movements of the IC region of TM6, which is one of the hallmarks of the class A GPCRs activation process (Zhou et al., 2019): the distance between the IC areas of TM3 and TM6 does not show remarkable differences between the three states despite the different values of θ (see also Figure S5), strengthening the previously observed different behaviour compared to class A GPCRs (Xu et al., 2022).

**Figure 2.**
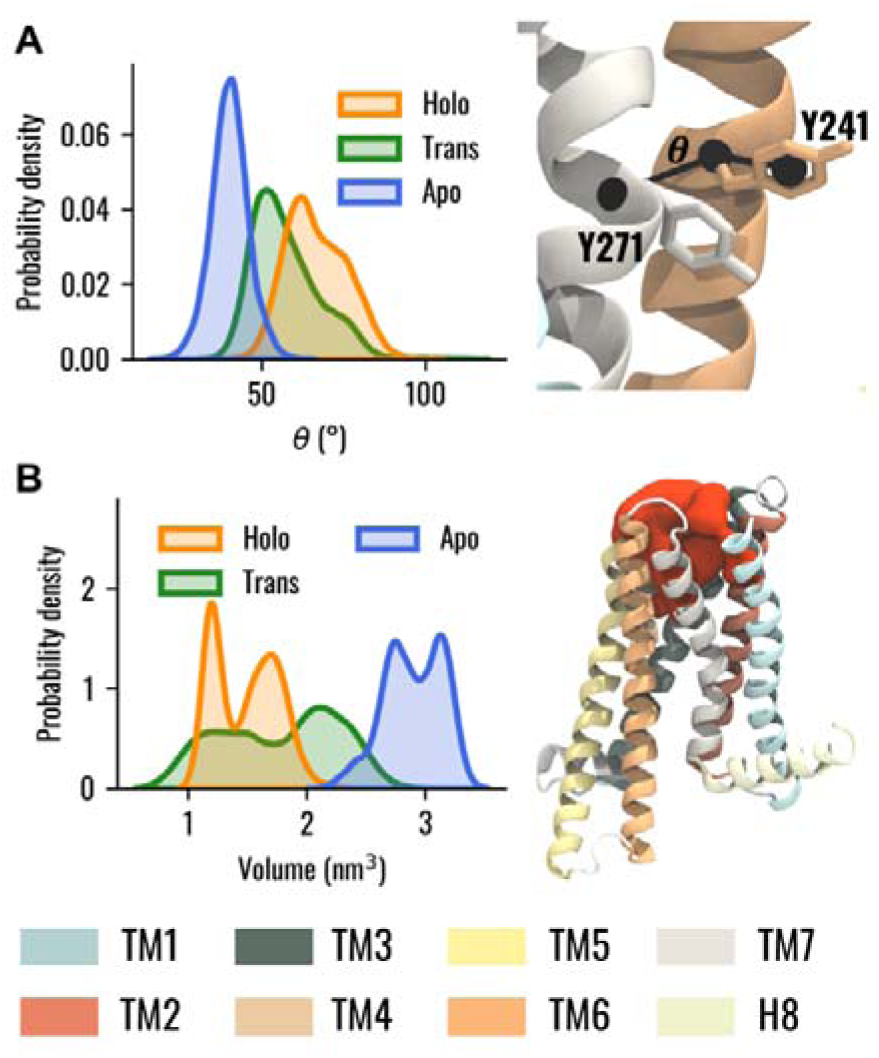
Conformational analysis of TAS2R46 in the three investigated states. (**A**) Probability distribution of the θ angle for Holo, Trans, and Apo states (left) and visual representation of angle θ definition (right). (**B**) Probability distribution of the binding pocket volume for the Holo, Trans, and Apo states (left) and visual representation of the binding pocket volume, coloured in red, for one selected frame of the Apo system (right). The ECL2 was removed for the sake of clarity.

Then, the volume of the TAS2R46 orthosteric binding pocket was evaluated in the three systems. It is worth mentioning that, in this analysis, the ECL2 was not considered to prevent any alterations in the estimation, as it is located within the investigated pocket in the Apo state. In this way, we evaluated only the change in the volume of the orthosteric binding pocket due to the rearrangements of the helices forming the ligand pocket, regardless of the initial location of the ECL2, which varies in the ligand-free or bound states. The results showed that the volume of the orthosteric pocket is higher in the absence of strychnine (Figure 2B). Interestingly, the Trans state displayed an intermediate behaviour as it is characterized by a distribution of the volume located between the ones of Apo and Holo states.

### 3.2 The ligand-bound state is characterized by an increased dynamical correlation

In this section, we assessed intra-structure correlations and examined how they were altered by the presence or absence of strychnine to identify major differences. The correlation between the different receptor regions in the three states was analyzed using the generalized correlation coefficients as described in the Material and Methods section. We evaluated the correlation between residues for the Holo, Trans, and Apo states, which showed remarkable differences in their dynamic behaviour. In the Holo system, high correlations were observed between the EC region of the receptor, whereas the IC regions of TM5 and especially TM6 were less correlated with the rest of the 7TM bundle (Figure 3A). Interestingly, the ICL3 displayed higher correlations than the neighbouring regions of TM5 and TM6, especially with the ECL1. Our results also showed that the removal of strychnine from the orthosteric binding pocket induced a remarkable and general loss of dynamic correlation (more bluish areas) within the 7TM bundle (Figure 3B). High correlations for the Trans state were observed only for the EC regions of TM1, TM6, and TM7, the ECL2, and, less markedly, the ECL1. On the other hand, the regions showing the lowest correlation with the 7TM bundle were the IC regions of TM3, TM6, and TM7. Finally, in the Apo state, the results highlighted remarkable correlations of the ECL2 with the EC regions of the receptor, while lower values could be observed for the IC regions (Figure 3C). To easily compare the receptor states, we also calculated the differences of the generalized correlation coefficient between the Trans and Holo states, i.e. Δr_MI_(Trans, Holo) = r_MI_(Trans) - r_MI_(Holo), and between the Apo and Holo states, i.e. Δr_MI_(Apo, Holo) = r_MI_(Apo) - r_MI_(Holo) (Figure 3D). Comparing the observed correlations for the three states, it was observed that, in general, the receptor evolved dynamically in a more correlated way in the presence of strychnine, whereas its absence was associated with the decorrelation of the IC regions. Moreover, it is worth noticing that the Trans state overall demonstrated lower structural correlation values compared to the other two. Furthermore, Figure 3D highlighted that the ICL3 and the IC portion of TM3 are less correlated with the rest of the 7TM bundle in Trans and Apo states than in the Holo system. Interestingly, for both Trans and Apo systems, the IC portion of TM6 is more correlated with the rest of the receptor compared to the Holo state. Therefore, the Apo and Trans systems behaved in the same way if compared to the Holo state.

**Figure 3:**
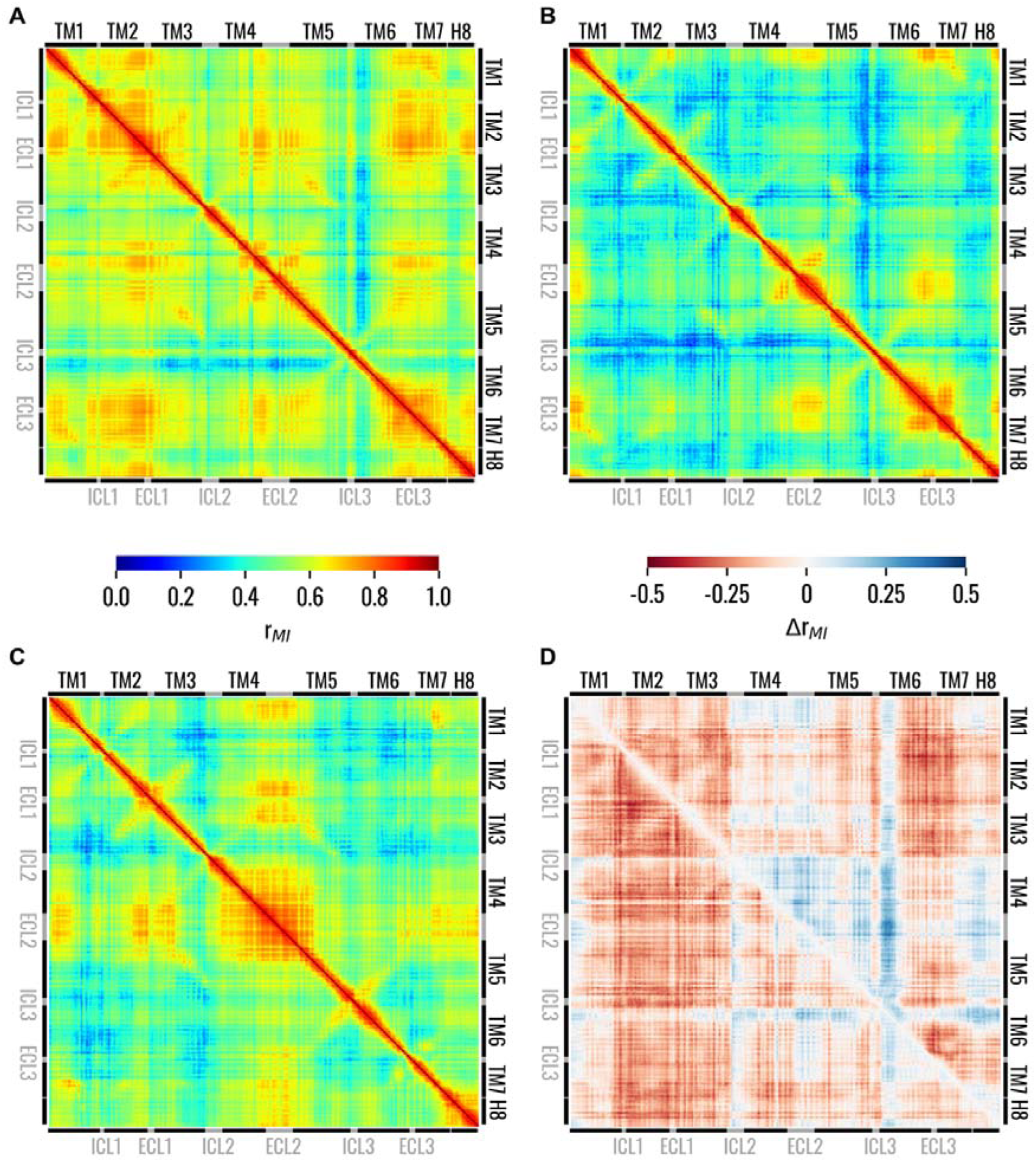
Matrices showing the generalized correlation coefficient between the residues of TAS2R46 in the (A) Holo, (B) Trans, and (C) Apo states. In panel (D), the differences of the generalized correlation coefficients are shown: Δr_MI_(Trans, Holo) = r_MI_(Trans) - r_MI_(Holo) in the lower triangle and Δr_MI_(Apo, Holo) = r_MI_(Apo) - r_MI_(Holo) in the upper triangle. Matrices on panels from A to C are coloured according to the left colorbar, while the matrix on panel D is colored according to the right colorbar.

### 3.3 Holo structural network is mediated by the TM3-TM6 connection

The dynamic networks of Holo, Trans, and Apo systems were initially analyzed by extracting information regarding node and edge centralities from the graph topology. Plotting the node and edge betweenness centrality versus the node and edge rank showed that both betweennesses had similar values in the three networks (Figure S6). The knees of these curves were identified, for each of the three systems and both node and edge betweennesses, and the lowest ones, belonging to the Holo state, were used as thresholds. Then, only nodes or edges with betweenness higher than the identified threshold were considered in the subsequent analysis. The distribution of node betweenness centralities in the different helices of the receptor was first considered, with specific attention to the upper tails of the distributions and outliers, which highlight residues of particular importance in the information flow inside the receptor. From this analysis, it was observed that, for the Holo state, the TM3 and TM6 were characterized by tails extending toward higher centralities, while other isolated residues were more central in other helices like TM1. On the other hand, for the Trans and Apo states, residues belonging to the TM3 were in general higher than the TM6 ones (p < 0.05 with Wilcoxon-signed-rank test), especially in the case of the Trans system (Figure 4A). Moreover, it could be observed that the tail of the distribution for TM4 extends toward higher values for Holo and Trans states compared to the Apo state. Finally, while the Holo and Apo systems showed similar ranges for TM2 and TM5 distributions, in these regions the Trans system was characterized by tails reaching higher centralities. Given that the betweenness centrality of nodes measures how influential a node is in the flow of information inside the network, these results highlight that the flow of information in TAS2R46 is mainly conveyed by the TM3 in the absence of strychnine, while by both TM3 and TM6 in the presence of the ligand. Similar information can be obtained from the visual inspection of the networks if the edges betweenness centrality is highlighted (Figure S7).

**Figure 4.**
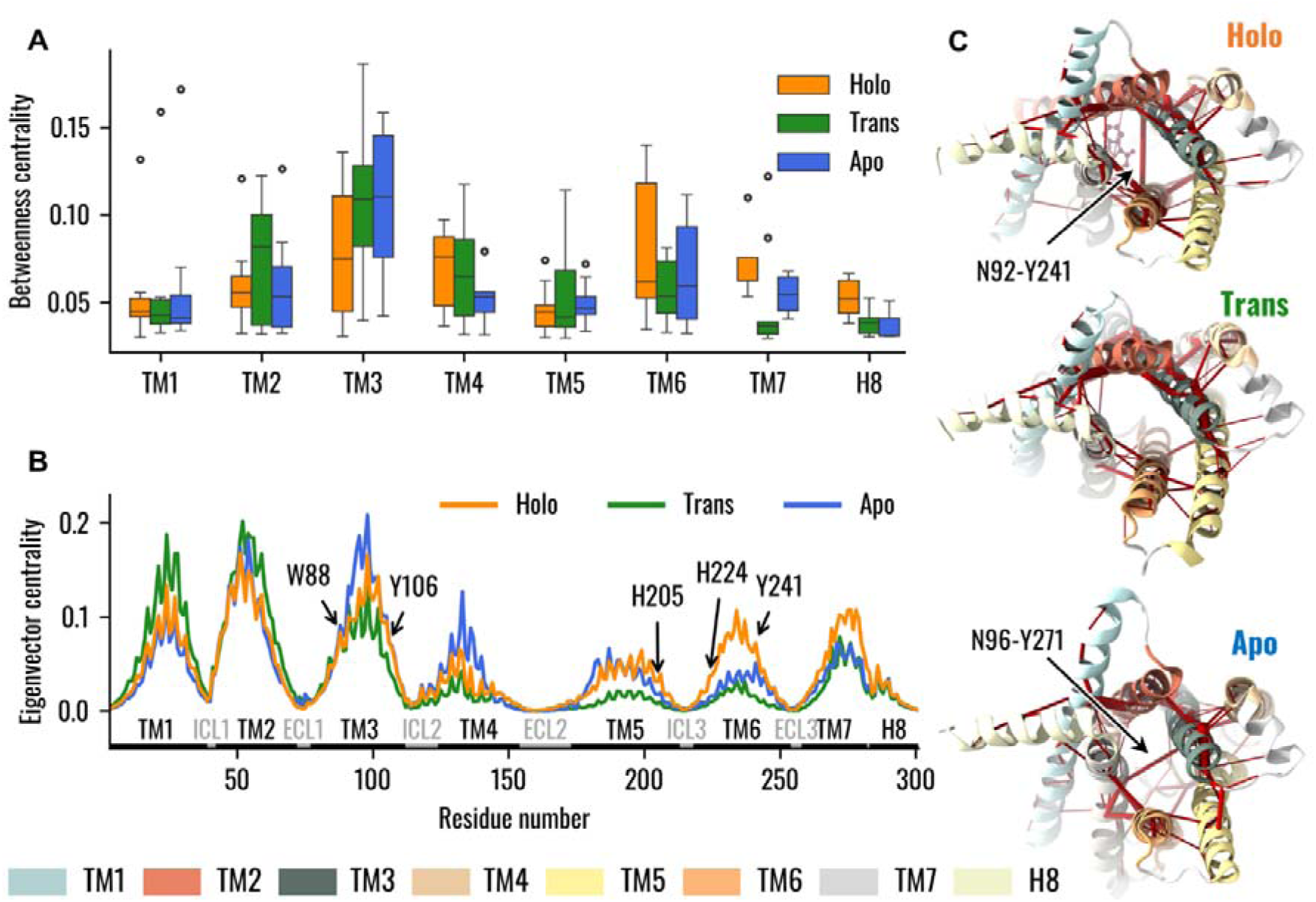
Dynamical Network Analysis of the Holo, Trans, and Apo systems. (**A**) Boxplots representing the distribution of node betweenness centrality, with nodes grouped into the respective TMs. Data emphasize that in the absence of the ligand TM3 assumed a primary role in information transfer, whereas in the presence of strychnine TM6 nodes also demonstrated high values of betweenness centrality. (**B**) Eigenvector centrality of TAS2R46 nodes. The TMs and loops are highlighted as well as residues pinpointed in previous experimental analysis. Results showed higher centrality of TM6 with strychnine, especially for H205^5.68^, H224^6.31^, and Y241^6.48^, and decreased centralities in the Trans state. (**C**) Visual representation of the dynamic networks, where the receptor is viewed from the IC region. The edges of the network are red cylinders with a radius proportional to the betweenness centrality of the edge.

The influence of the single residues on the overall network was also analyzed in terms of eigenvector centrality. The results showed similar values for the EC and IC portions of the TM3 in the Apo and Holo systems, while the TM6 was characterized by remarkably higher centrality in the presence of strychnine (Figure 4B). In particular, while W88^3.32^, involved in strychnine binding, and Y106^3.50^, involved in G-protein interaction at the TM3 level (Xu et al., 2022), displayed similar centralities in Holo and Apo states, H205^5.68^, H224^6.31^, and Y241^6.48^ were more central in the presence of the ligand. Moreover, except for W88^3.32^, all mentioned residues showed lower centralities in the Trans systems. This is particularly interesting as H205^5.68^ and H224^6.31^ are involved in G-protein interaction at the TM5 and TM6 levels, respectively (Xu et al., 2022). Moreover, as discussed before, Y241^6.48^ has to have a pivotal role in TAS2R46 activation.

Finally, we focused on the difference, between the three systems under analysis, in terms of connections between TMs. Interestingly, the results showed that the main difference is relative to the TM3 (Figure S8): whereas in the presence of the ligand, there is a connection between TM3 and TM6 mediated by the interaction between N92^3.36^ and Y241^6.48^, in the ligand-free state TM3 is connected to TM7 through the N96^3.40^-Y271^7.45^ interaction, and in the Trans state there is no connection between TM3 and TM6 or TM7 (Figure 4C and see also Figure S8).

To investigate the effects of the highlighted changes in the dynamic network properties on the structural communication inside TAS2R46, we computed the optimal paths linking strychnine binding residues, in particular W88^3.32^ and E265^7.39^, to H224^6.31^ and Y106^3.50^, as in a previous study they were identified as G-protein binding residues (Xu et al., 2022). In line with the analysis of the network topology (Figure S8), in all systems the communication between W88^3.32^ and H224^6.31^ is driven by the TM3. On the other hand, differences could be observed when considering the W88^3.32^-H224^6.31^ and E265^7.39^-Y106^3.50^ paths (Figure 5, see also Table S1). In the presence of strychnine, both paths involve the edge between N92^3.36^ and Y241^6.48^, which creates a bridge between TM3 and TM6 through which the information between the EC and IC regions is conveyed. In the Apo state, instead, the path from W88^3.32^ to H224^6.31^ reaches the IC region of the receptor mainly through the TM3, then passes to the TM5, and finally to H224^6.31^. Interestingly, the same path was observed in the Trans state, with the only difference being that the path reaches TM6 directly at H224^6.31^ residue, instead of A227^6.34^ as in the Apo state. On the other side, the connection between E265^7.39^ and Y106^3.50^ for the Apo system is mediated by the Y271^7.45^-N96^3.40^ interaction linking TM3 and TM7, whereas in the Trans and Holo systems, the paths are mediated by the TM2 and TM6, respectively (Figure 5, see also Table S1). Therefore, the change in the network topology is reflected in remarkable differences in the communication between the EC and IC regions.

**Figure 5.**
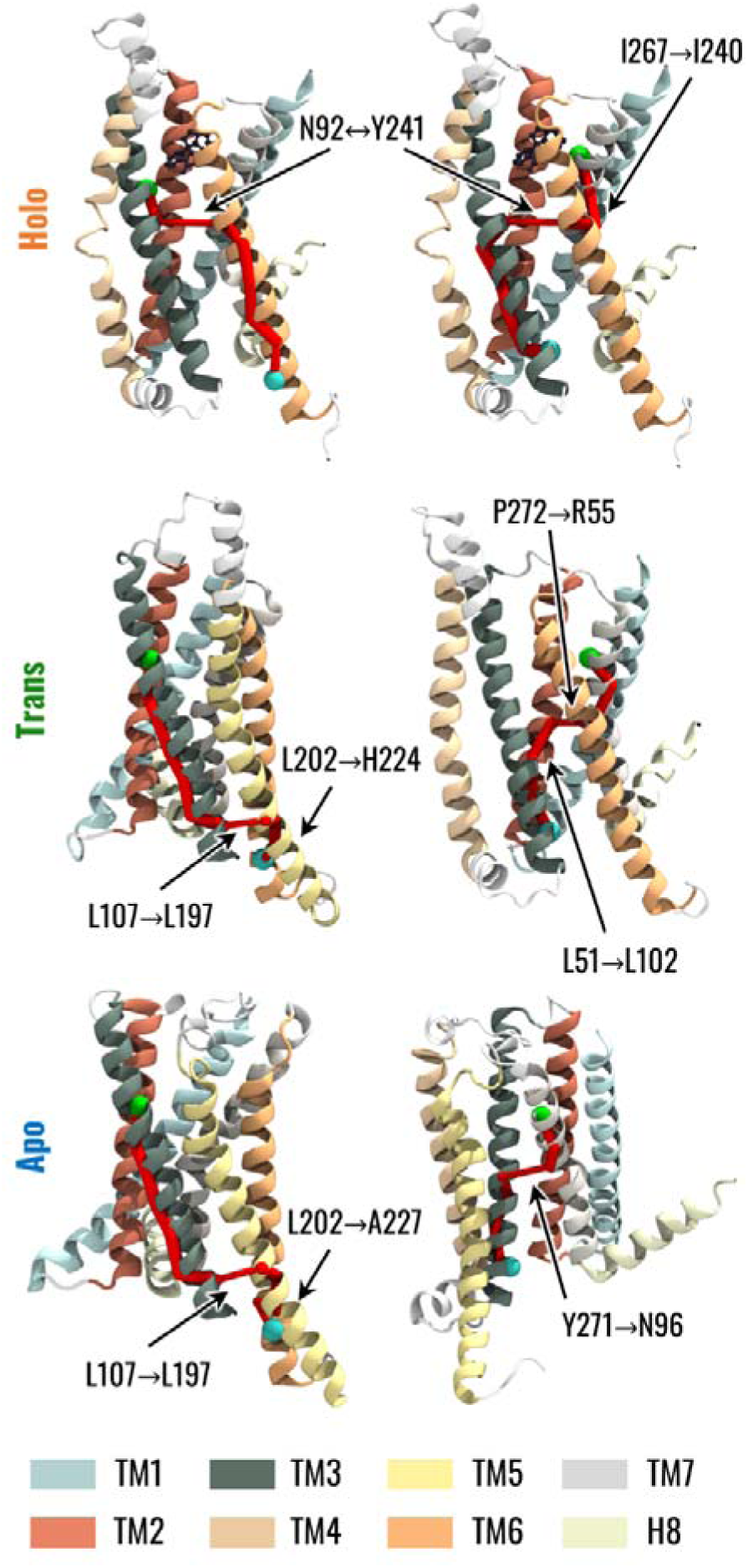
Visual representation of the optimal path connecting strychnine- and G protein-binding residues. The left column represents the communication path between W88^3.32^ in TM3, highlighted as a green sphere, and H224^6.31^ in TM6, represented as a cyan sphere. The right column represents the communication path between E265^7.39^ in TM7, highlighted as a green sphere, and Y106^3.50^ in TM3, represented as a cyan sphere. The edges in the paths are represented by cylinders whose radius is proportional to the generalized correlation coefficient between residues.

## 4 Discussion

TAS2Rs constitute the molecular basis of bitter taste perception. Concurrently, there is mounting evidence that various extra-oral tissues express bitter taste receptors, and their activation elicits diverse signals and cellular responses essential for metabolism and homeostasis (Dotson et al., 2008; Liggett, 2014; Pan et al., 2017). Extra-oral TAS2Rs are associated with several diseases, and they could represent a promising target for pharmacological intervention. However, the lack of experimental structures of TAS2Rs represented an important hurdle for understanding the mechanisms underlying bitter receptor activation. The recent release of the experimental structures of TAS2R46 both coupled with strychnine and in a ligand-free state (Xu et al., 2022) paves the way for the structural characterization of this receptor and for the definition of specific dynamic hallmarks underlying the receptor state. Strychnine is of important interest as a ligand because it is experimentally known to target not only TAS2R46 but also other bitter taste receptors, such as TAS2R10, and even other proteins (Born et al., 2013; Brockhoff et al., 2010; Jensen et al., 2006; Sandal et al., 2015; Xue et al., 2018). In this context, this work aims to contribute to the overall understanding of the molecular mechanisms related to the activation of TAS2Rs. In detail, the conformational and dynamical features of the receptor in the presence and absence of strychnine were considered and then the dynamic network analysis was employed to investigate how such features could be related to the allosteric activation, which implies the transfer of information from the EC region to the IC region. Indeed, network analysis of protein dynamics has already proven their potential for analyzing the structural communication in macromolecular structures, including GPCRs (Bertalan et al., 2020; Bondar, 2022; Fanelli et al., 2016; Melo et al., 2020; Sullivan et al., 2020).

Initially, conformational features typical of class A GPCRs or highlighted by previous experimental studies were considered. Regarding the interaction between strychnine and TAS2R46, the PLIP analysis showed that the initial interactions with W88^3.32^ and E265^7.39^ remained stable during the MD simulation, and at the same time pinpointed a stable hydrophobic interaction with Y85^3.29^ (Figure S4). It is worth mentioning that these residues have been highlighted in previous literature to be involved in contacts with strychnine in TAS2R46 (Sandal et al., 2015; Xue et al., 2018) or agonist selectivity in other TAS2Rs, including TAS2R38 (Marchiori et al., 2013), TAS2R10 (Born et al., 2013), TAS2R16 (Sakurai et al., 2010), TAS2R43 (Brockhoff et al., 2010; Pronin, 2004), TAS2R44 (Brockhoff et al., 2010; Pronin, 2004), TAS2R47 (Pronin, 2004), TAS2R1 (Singh et al., 2011), and TAS2R4 (Pydi et al., 2012). Additionally, residue W88^3.32^ is involved in the activations of TAS2R43 and TAS2R30 (Pronin, 2004), whereas residue E265^7.39^ was previously reported to be involved in a salt-bridge interaction with strychnine (Xue et al., 2018) and its mutation implicates a reduced responsiveness to the compound (Brockhoff et al., 2010). However, while residue W88 is highly conserved among TAS2Rs (84%), residues E265 and Y85 show a lower conservation (32% and 24%, respectively), suggesting a different impact on receptor selectivity. Furthermore, residues in positions 3.32 and 7.39 are known to be involved in the binding of diverse ligands in class A GPCRs (Brockhoff et al., 2010; Venkatakrishnan et al., 2013).

Nevertheless, our analysis corroborated the hypothesis from previous findings regarding the lack of class A GPCR conformational activation hallmarks (Bock and Bermudez, 2021; Ibrahim et al., 2019; Zhou et al., 2019). Specifically, our simulations did not reveal a remarkable outward movement of TM6 (Figure S5) or an increase in the A^100^ index (Figure S9), indicating that these markers may not adequately describe the activation mechanism of TAS2Rs. These observations strengthen the recent new classification of TAS2Rs into the class T of GPCRs (Pándy-Szekeres et al., 2023), characterized by distinctive features different from class A (Di Pizio et al., 2016; Topin et al., 2021). On the contrary, as was observed for the class A GPCRs activation process (Dalton et al., 2015), in the TAS2R46 strychnine-bound state the volume of the orthosteric binding pocket is lower than the one in the Apo state and the removal of strychnine from the Holo system was associated with an increase in the pocket volume (Figure 2B). Interestingly, the Trans state demonstrated intermediate binding pocket volume values between those of the Apo and Holo states.

The overall dynamic of TAS2R46 was influenced by the presence or absence of strychnine in the orthosteric binding site. This was highlighted by the analysis of the intra-receptor correlations, which showed higher overall correlation values for the Holo state, while both the Trans and the Apo states were characterized by lower correlations (Figure 3). In particular, in Trans and Apo states, reduced correlation for the IC region of TM3 and for the ICL3, which are important regions for G-protein binding, was observed. Therefore, the Apo and Trans systems were characterized by similar behaviour in terms of intra-receptor correlations if compared to the Holo state. The main difference between the Apo and Trans states could be linked to the different conformation of the ECL2. Indeed, in the initial state, the ECL2 is placed within the orthosteric binding site in the Apo state and outside in the Trans state. During the MD simulation, the ECL2 showed remarkable correlations with other regions of the receptor only for the Apo structure. Furthermore, it is noteworthy that the removal of strychnine in the Trans state resulted in a globally less correlated state if compared to the Apo and Holo states. Therefore, the whole receptor in the Trans state seemed to evolve in time in a more decorrelated way.

The different conformational and dynamical features of the receptor in the three states were then related to different behaviours of the receptor in terms of structural communication using the dynamic network analysis. The betweenness centrality pointed out the importance of TM3 in the information transfer from the EC to the IC region of the receptor independently of its state (Figure 4). This finding is of particular interest given the pivotal role of TM3 in class A GPCRs, serving as a structural and functional hub responsible for maintaining the receptor scaffold in both active and inactive states (Venkatakrishnan et al., 2013). Thus, these results support the hypothesis that TM3 represents a central and critical structural element that defines the overall structure of both class A GPCRs and TAS2Rs in a similar manner. On the other hand, the TM6 was characterized by increased importance and influence inside the network in the presence of strychnine (Holo state) (Figure 4A,B). This information is particularly relevant as it highlights that high values of the centrality of TM6 in the presence of strychnine are related to the above-mentioned connection to TM3 through the interaction between Y241^6.48^ and N92^3.36^ (Figure 4C). Moreover, it is worth noting that the central role of TM6 in the activation process has also been highlighted in previous literature related to class A GPCRs (Trzaskowski et al., 2012; Venkatakrishnan et al., 2013; Zhou et al., 2019). Regarding the network connecting the TMs, the dynamic network analysis revealed distinct correlations for the Holo, Trans, and Apo structures (Figure 4C and see also Figure S8). Specifically, the Holo structure exhibited a high correlation between TM3 and TM6, while the Apo structure showed an edge connecting TM3 and TM7. Interestingly, the dynamic network of the Trans structure did not show any edges connecting TM3-TM6 or TM3-TM7. These results once more indicate a possible remarkable distinction between class A GPCRs and TAS2Rs. Specifically, following activation in class A receptors, the outward movement of the TM6 helix (not relevant for TAS2Rs as noted above) results in a reduction in contacts between the TM3 and TM6 helices and the formation of contacts between the TM3 and TM7 helices (Venkatakrishnan et al., 2016; Zhou et al., 2019), thus showing a different trend to that observed during the dynamics of TAS2R46.

We specifically focused our attention on the residue at position 6.48, which is considered crucial in the activation process of class A GPCRs (Tokmakova et al., 2023) and has been suggested to be involved in the “toggle switch” mechanism in previous work related to the experimental resolution of the TAS2R46 structure (Xu et al., 2022). Among the side chain rearrangements during the MD simulations, we quantified the rotation of Y241^6.48^, which has been highlighted in the experimental structures (Xu et al., 2022). The MD simulations confirm that the side chain of Y241^6.48^ points towards the centre of the 7TM bundle in the presence of strychnine, forming an interaction with N92^3.36^, and towards the TM7 in the Apo states (Figure 2A). The interaction of Y241^6.48^ and N92^3.36^ was also reported in previous literature regarding TAS2R46 bound to strychnine (Sandal et al., 2015). Additionally, Y241^6.48^ and N92^3.36^ demonstrate high conservation among TAS2Rs (64% and 84%, respectively) and the mutations of such residues lead to reduced sensitivity and response levels, also in other TAS2Rs, confirming their importance in the activation process (Pronin, 2004; Sandal et al., 2015). Interestingly, the removal of strychnine altered the conformation of Y241^6.48^, which showed a tendency to rotate towards the TM7, placing the Trans state between the others. Similar to our analysis, recent literature has also pointed out a similar interaction between residues Y^6.48^ and N^3.36^ through a hydrogen bond in the predicted active state of TAS2R14 (Tokmakova et al., 2023). In contrast, in class A GPCRs, the side chain of the highly conserved residue W^6.48^ alternates between Gauche+ and trans conformations for the active and inactive states, respectively (Tokmakova et al., 2023). This highlights a distinct difference in behaviour between class A receptors and TAS2Rs. Specifically, residue W^6.48^ in class A GPCRs is able to create a bridge between TM6 and TM3 through an interaction with residue in position 3.36 in the inactive state (Trzaskowski et al., 2012; Venkatakrishnan et al., 2013).

Concerning the dynamic network analysis involving residue Y241^6.48^, we pointed out the formation in the Apo state of a different edge connecting TM3 and TM7 due to the rotation of residue Y241^6.48^ towards the TM7. At the same time, the influence of TM6 in the network is reduced. On the other hand, the intermediate localization of the Y241^6.48^ side chain for the Trans system did not allow the connection of TM3 either with TM6 or TM7, again placing the behaviour of the Trans structure between Holo and Apo states. The pivotal position of Y241^6.48^ was also confirmed by its implication in the optimal paths linking strychnine binding residues (W88^3.32^ and E265^7.39^) to previously identified G-protein binding residues (H224^6.31^ and Y106^3.50^) (Xu et al., 2022), as represented in Figure 5 and reported in Table S1. It is worth mentioning that residues H224^6.31^ and Y106^3.50^ are highly conserved among TAS2Rs (88% and 92%, respectively), underscoring their importance for the majority of the bitter taste receptors. In all systems (Apo, Trans and Holo) the communication between W88^3.32^ and H224^6.31^ is mediated by TM3, whereas the connection between E265^7.39^ and Y106^3.50^ is facilitated by TM6 only in the Holo state. In the presence of strychnine (Holo state), both paths involve the edge between N92^3.36^ and Y241^6.48^, which creates a direct bridge between TM3 and TM6. On the contrary, in the absence of the ligand (Apo and Trans states) the path from W88^3.32^ to H224^6.31^ involves the creation of an edge connecting TM3 and TM5, before reaching TM6. On the other hand, the connection between E265^7.39^ and Y106^3.50^ is mediated by a direct connection linking TM3 and TM7 (Y271^7.45^-N96^3.40^) in the Apo system, whereas this path involves the formation of the network between TM7-TM2-TM3 in the Trans state. In summary, it is noteworthy that the dynamic network analysis revealed that the mechanical information is transmitted from the orthosteric binding site to the intracellular (IC) domain via a pathway involving TM3-TM6 in the presence of strychnine, directly implicating residue Y241^6.48^ in the network edge. Additionally, the removal of the ligand from the binding site caused the Trans structure to form pathways resembling those of the Apo state, connecting residues W88^3.32^ to H224^6.31^.

In conclusion, this study suggests how the “toggle-switch” represented by Y241^6.48^ side chain rotation could induce TAS2R46 allosteric activation without evident conformational changes in the IC region of the receptor. The rotation of Y241 in TM6 towards the centre of the 7TM bundle allows for the formation of an interaction with N92^3.36^ in TM3, which forms a bridge through which the mechanical information can be transferred between the two helices whose IC regions are involved in G-protein binding. This is also associated with different dynamical behaviour of the receptor, which, in the presence of strychnine, is characterized by higher correlations between the IC and EC regions. In this way, the allosteric network generated by ligand binding is directly transferred to the G-protein binding sites, possibly inducing G-protein activation.

It is essential to emphasize that the present study includes some limitations to be addressed in future works. First, the simulated structures did not include the G-protein within the models and therefore cannot directly quantify the effects of the presence or absence of the bitter agonist on the G-protein. Moreover, the TAS2R46 structures were embedded into a homogeneous POPC bilayer, despite it is known that the membrane composition can modulate GPCRs function, stability, and signalling (Gimpl, 2016; Sengupta et al., 2018). Therefore, further studies will be fundamental to expand the results of the present work towards a complete characterization of the TAS2R46 structure and dynamics. However, this study represents an important advance toward the comprehension of TAS2R activation mechanisms, highlighting specific structural dynamic hallmarks of TAS2R46 and pinpointing major differences compared to class A GPCRs.

## 5 Conclusion

The present study investigates molecular mechanisms and features governing the activation of a G protein-coupled receptor, namely the human TAS2R46 bitter taste receptor. Through a computational approach involving molecular dynamics simulations and network-based analysis, we provided key insights into conformational properties, global intra-structural correlations and allosteric networks associated with TAS2R46 bound or unbound to a bitter agonist. The results highlighted that TM3 and TM6 are the main helices involved in the allosteric network when the receptor is bound to strychnine, while TM6 reduces its influence when strychnine is absent. Moreover, molecular simulations confirmed previous evidence regarding the importance of Y241^6.48^ side-chain localization in the activation process and provided additional insights into the effect of such localization on the allosteric network of the receptor. Lastly, we emphasised that the presence of the bitter agonist increased the overall intra-structural correlations, and the removal of the ligand resulted in a loss of these correlations. The results of the present work also highlighted the similarities and differences between class A GPCRs and TAS2Rs, supporting the hypothesis that bitter taste receptors should be classified into a distinct family, characterised by unique structural and dynamic features. Hence, the proposed methodology could potentially be expanded to characterise the molecular mechanisms and allosteric networks of other TAS2Rs, aiming to identify both common and specific features in the activation process, which may vary given the wide variability in this fascinating class of receptors.

In conclusion, this study has unveiled crucial molecular features and essential allosteric networks involved in the TAS2R46 function. These findings enhance our comprehension of the intricate activation processes within the bitter taste receptor family and the present approach holds promise for characterizing similar protein targets in future studies. Given the obtained results, further exploration of the activation process of TAS2R46 in the presence of various agonists is warranted to discern whether different agonists trigger similar signal transduction mechanisms. Therefore, this work serves as a pivotal starting point for a profound understanding of the precise mechanisms governing TAS2R activation, paving the way for the rational design of small compounds targeting oral or extra-oral TAS2Rs.

## 6 Acknowledgements

The present work has been developed as part of the VIRTUOUS project, funded by the European Union’s Horizon 2020 research and innovation program under the Marie Sklodowska-Curie-RISE Grant Agreement No. 872181 (https://www.virtuoush2020.com/).

## 7 Author contributions

**Marco Cannariato**: Conceptualization; Data curation; Formal analysis; Methodology; Software; Validation; Visualization; Writing - original draft; Writing - review & editing. **Riccardo Fanunza**: Conceptualization; Data curation; Formal analysis; Investigation; Validation; Visualization; Writing - review & editing. **Eric A. Zizzi**: Supervision; Writing - review & editing; **Marcello Miceli**: Supervision; Visualization; Writing - review & editing; **Giacomo Di Benedetto**: Supervision; Writing - review & editing; **Marco Agostino Deriu**: Conceptualization; Data curation; Funding acquisition; Project administration; Resources; Supervision; Validation; Visualization; Writing - review & editing; **Lorenzo Pallante**: Conceptualization; Data curation; Methodology; Software; Supervision; Validation; Visualization; Writing - review & editing.

## 8 Data and Software Availability

All the files necessary to reproduce the simulations and the Python scripts for the analysis are accessible at the website https://github.com/lorenzopallante/TAS2R46.

## 9 Declaration of interests

The authors declare that no financial interest or conflicts of interest exist.

## Notes

https://github.com/lorenzopallante/TAS2R46

### Competing Interest Statement

The authors have declared no competing interest.

### Summary of Updates

Main changes in the Introduction and Discussion sections.

