## Supplementary Information for "Exploring TAS2R46 Biomechanics through Molecular Dynamics and Network Analysis"

[b]7HC s.r.l., 00198, Rome, Italy

### *Supplementary Material*

### 1 Supplementary Methods

As TAS2Rs have been considered to be class A GPCR until human TAS2R46 structure was resolved, the $A^{100}$ activity indicator, specifically developed to describe class A GPCR activation, was computed as reported in previous literature (Calderón et al., 2023; Ibrahim et al., 2019). In particular, the $A^{100}$ is computed as:

$$A^{100} = -14.43\ R^{I27,W281} - 7.62\ R^{R55,H93} + 9.11\ R^{L98,S129} - 6.32\ R^{C203,H228} - 5.22\ R^{S252,F261} + 278.88$$

where $R$ stands for the distance between the alpha carbons of the residues given in the superscript.

Moreover, as the outward movement of the TM6 IC region is one of the hallmarks of the class A GPCRs activation process (Zhou et al., 2019), the same conformational change was characterized. In particular, the distance between the IC areas of TM3 and TM6 in the three states was computed as the distance between the centre of mass of residues 104 to 100 (TM3) and the centre of mass of residues 218 to 224 (TM6).

### 2 Supplementary Figures

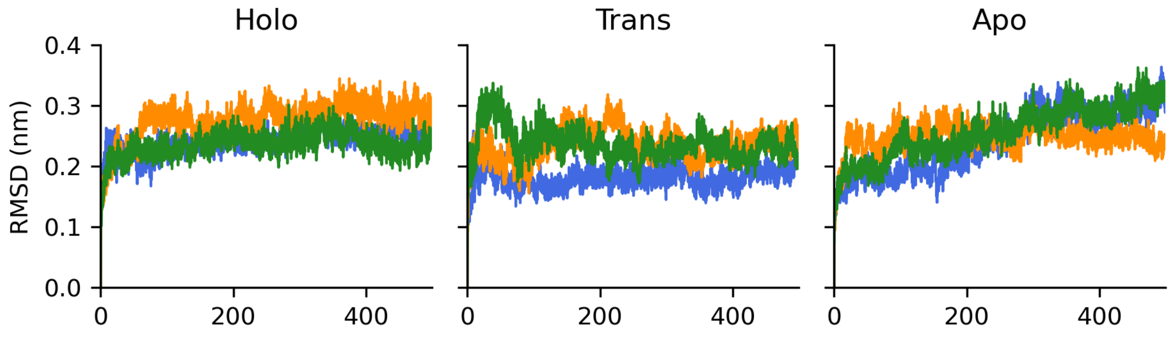


Figure S1. RMSD of backbone atoms with respect to the initial conformation of the MD simulations for the Holo, Trans, and Apo states. The three replicas are displayed in different colours.

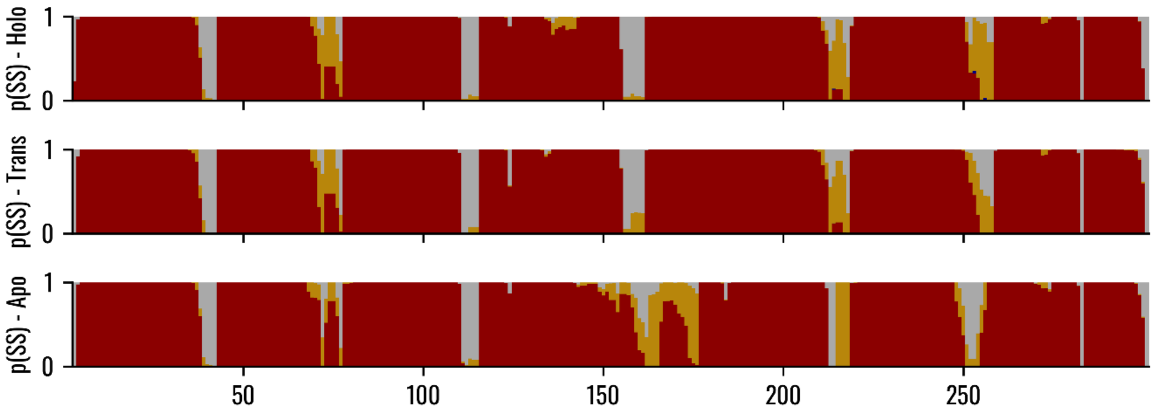


Figure S2. Probability of each TAS2R46 residue to be involved in the formation of a secondary structure. Helices are reported in dark red, beta sheets in dark blue, turns in dark yellow, and coils in grey.

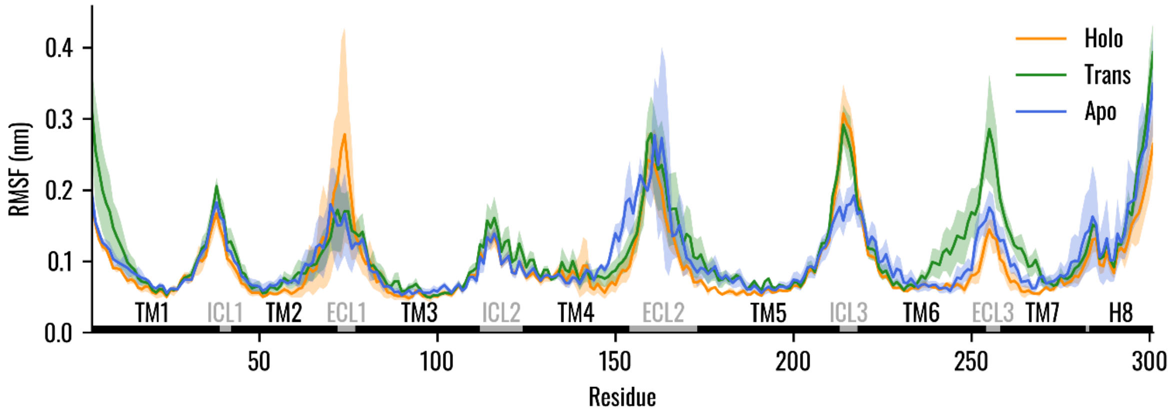


Figure S3. RMSF of alpha carbons computed on the equilibrium trajectories (last 400 ns). The mean value across the three replicas is represented with a continuous line, whereas the shaded regions represent the standard deviation from the mean.

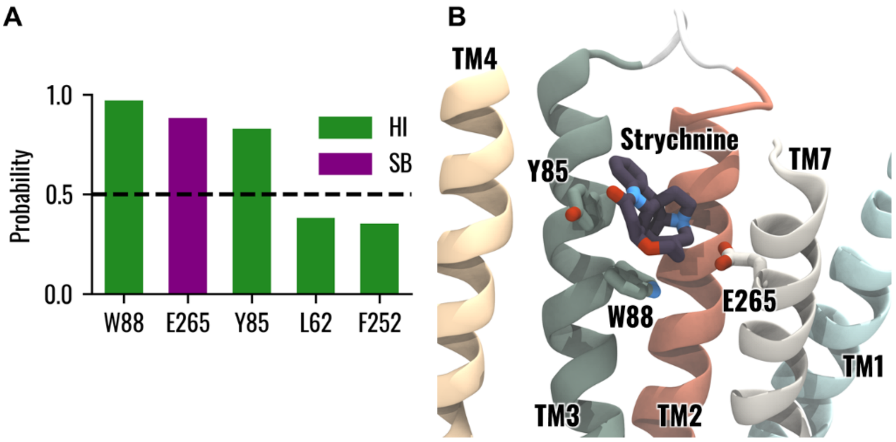


Figure S4. (**A**) Probability of TAS2R46 residues to form an interaction with strychnine, as identified by PLIP software. The bars are coloured according to the specific type of interaction, i.e. HI=hydrophobic interaction (green), and SB=salt bridges (purple). (**B**) Visual representation of strychnine in TAS2R46 binding pocket. Residues forming interactions with a probability greater than 0.5 with Strychnine are shown.

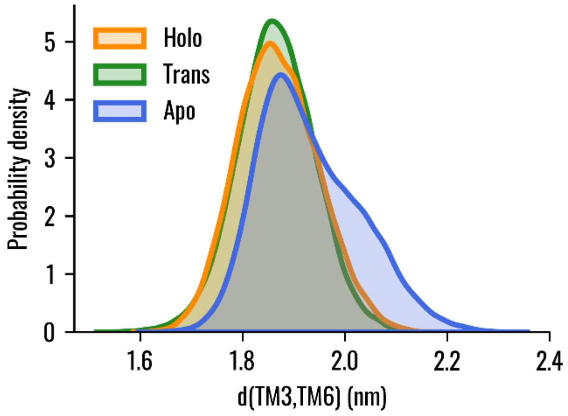


Figure S5. Distribution of the distance between the intracellular areas of TM3 and TM6 in the three states, which has been used to characterize possible outward movement of TM6. The centre of mass of residues 104 to 100 has been considered to define the intracellular portion of TM3, whereas the centre of mass of residues 218 to 224 to define the intracellular area of TM6.

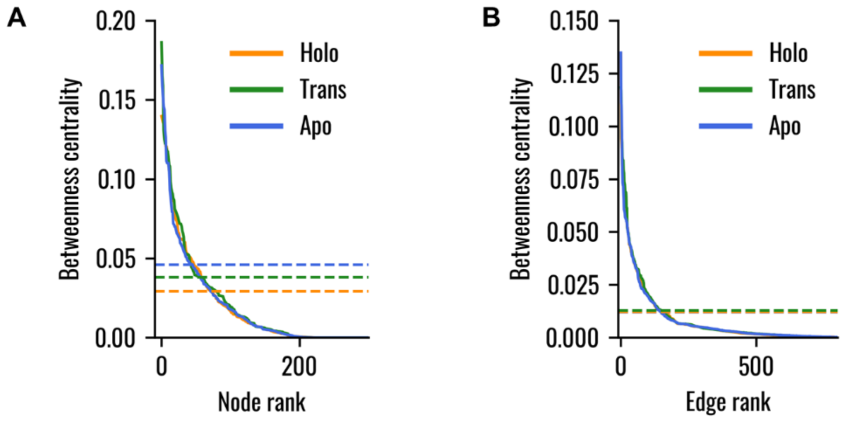


Figure S6. Ranked betweenness centrality for the (A) nodes and (B) edges forming the dynamic networks of the Holo, Trans, and Apo systems. The horizontal dashed lines represent the value at which a knee was identified in the three systems. The identified threshold was the lowest knee identified.

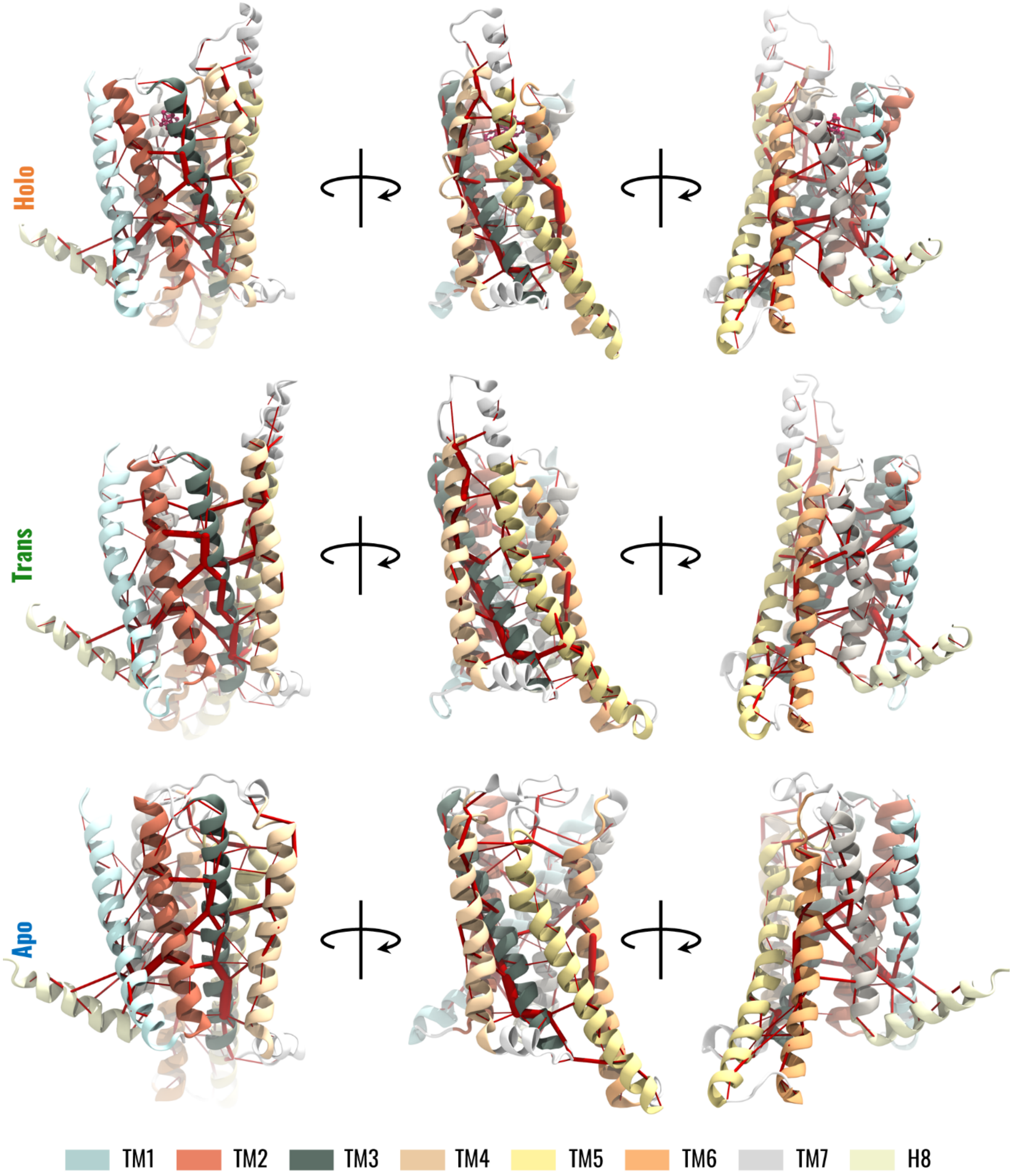


Figure S7. Visual representation of the dynamic network for the three states of the receptor. The edges of the network are red cylinders with a radius proportional to the betweenness centrality of the edge.

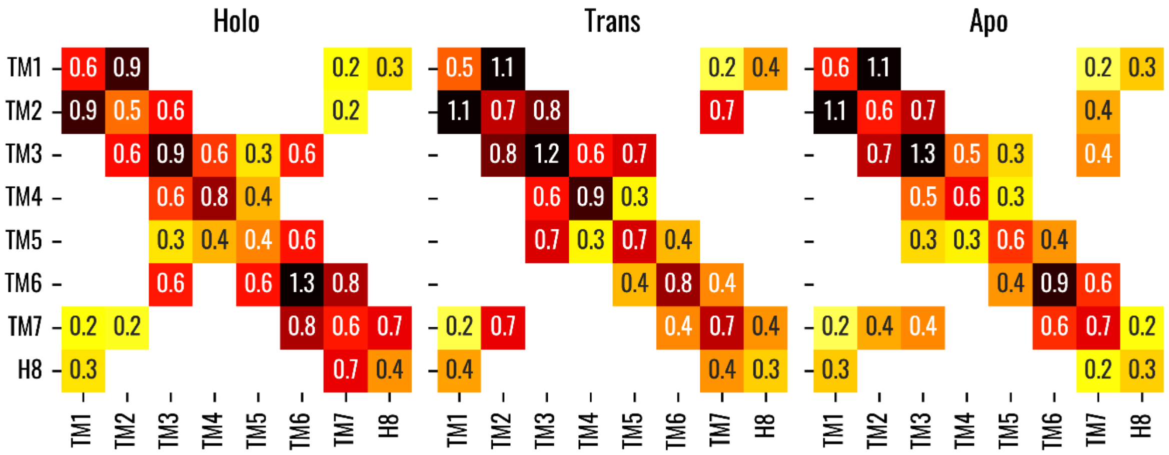


Figure S8. Maximum betweenness of the edges linking different TM helices. Values represent the maximum betweenness, which has been multiplied by 10 for a better representation. White regions correspond to the absence of edges between the corresponding TM helices.

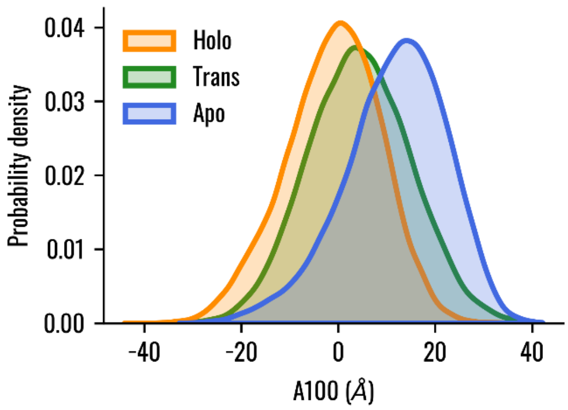


Figure S9. Distribution of the A$^{100}$ indicator in the Holo, Trans, and Apo states. It is worth mentioning that the change in the A100 index across the three systems demonstrated a different behaviour compared to what was proposed for class A GPCRs (Ibrahim et al., 2019).

Table S1. Summary of the optimal path connecting strychnine- and G protein-binding residues (visually represented in Figure 5 in the main text).

| | **W88$^{3.32}$ – H224$^{6.31}$** | **E265$^{7.39}$ – Y106$^{3.50}$** |
|---|---|---|
| **APO** | **TM3 – TM5 – TM6**<br>[L107$^{3.51}$ → L197$^{5.60}$]<br>[L202$^{5.65}$ → A227$^{6.34}$] | **TM7 – TM3**<br>[Y271$^{7.45}$ – N96$^{3.40}$] |
| **TRANS** | **TM3 – TM5 – TM6**<br>[L107$^{3.51}$ → L197$^{5.60}$]<br>[L202$^{5.65}$ → H224$^{6.31}$] | **TM7 – TM2 – TM3**<br>[P272$^{7.46}$ – R55$^{2.50}$]<br>[L51$^{2.46}$ – L102$^{3.46}$] |
| **HOLO** | **TM3 – TM6**<br>[N92$^{3.36}$ – Y241$^{6.48}$] | **TM7 – TM6 – TM3**<br>[I267$^{7.41}$ – I240$^{6.47}$]<br>[Y241$^{6.48}$ – N92$^{3.36}$] |
